# Rigorous surveillance is necessary for high confidence in end-of-outbreak declarations for Ebola and other infectious diseases

**DOI:** 10.1101/485821

**Authors:** Robin N. Thompson, Oliver W. Morgan, Katri Jalava

**Affiliations:** University of Oxford, Oxford, UK (RNT); World Health Organization, Geneva, Switzerland (OWM); University of Helsinki, Helsinki, Finland (KJ)

**Keywords:** Ebola virus disease, End-of-outbreak declarations, World Health Organization, Outbreak forecasting, Surveillance

## Abstract

The World Health Organization considers an Ebola outbreak to have ended once 42 days have passed since the last possible exposure to a confirmed case. Benefits of a quick end-of-outbreak declaration, such as reductions in trade/travel restrictions, must be balanced against the chance of flare-ups from undetected residual cases. We show how epidemiological modelling can be used to estimate the surveillance level required for decision-makers to be confident that an outbreak is over. Results from a simple model characterising an Ebola outbreak suggest that a surveillance sensitivity (i.e. case reporting percentage) of 79% is necessary for 95% confidence that an outbreak is over after 42 days without symptomatic cases. With weaker surveillance, unrecognised transmission may still occur: if the surveillance sensitivity is only 40%, then 62 days must be waited for 95% certainty. By quantifying the certainty in end-of-outbreak declarations, public health decision-makers can plan and communicate more effectively.

## Introduction

The 2018 Ebola outbreak in Equateur Province, Democratic Republic of the Congo (DRC), was brought under control following 54 cases between 5^th^ April and 2^nd^ June (*1*). Another unconnected outbreak was declared in DRC on 1^st^ August 2018, and that outbreak is still in progress. It is about to become the second largest in history, with 421 probable and confirmed cases as of 26^th^ November 2018 (*2*). Increasingly, decision-makers use forecasts generated using mathematical models to guide control measures when outbreaks are ongoing (*3,4*). However, less attention has been directed towards using mathematical modelling to inform decision-making at the ends of outbreaks (*5*).

Determining when an outbreak of any infectious disease is over is important for decision-makers, as they need to choose when to relax control measures, scale-back the deployment of personnel and resources, adjust communication messages to the public, and re-establish confidence in commercial sectors such as agriculture and tourism. However, the difficulty of end-of-outbreak decision-making was illustrated during the 2013-16 Ebola epidemic, when the World Health Organization (WHO) declared Liberia disease-free four times only to have new cases detected after the first three declarations (Fig 1A). This raises an important question: how confident can public health decision-makers be when declaring an outbreak over?

**Fig. 1.**
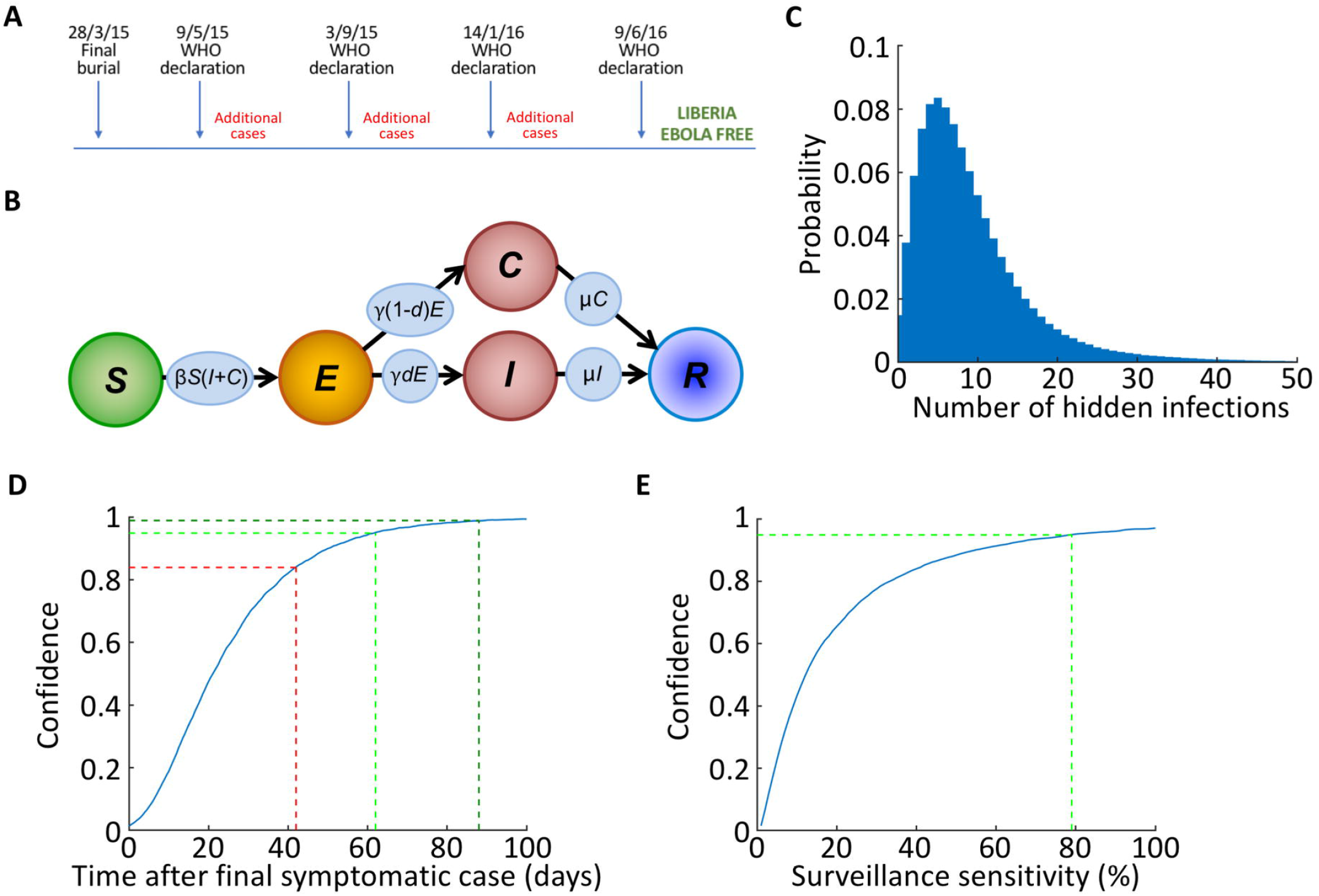
The confidence in end-of-outbreak declarations following the apparent end of an Ebola outbreak. A) Schematic showing the sequence of events in Liberia at the end of the 2013-16 Ebola epidemic, in which the outbreak was incorrectly declared over three times. B) Schematic of the compartmental model used in our analyses. C) The number of hidden cases (*E* or *C*) remaining in simulated Ebola outbreaks at the first timepoint at which the number of symptomatic cases (*I*) reaches zero. D) The confidence in end-of-outbreak declarations (i.e. the probability that no undetected infections (*E* or *C*) remain in the population), for different time periods after removal of the “final” symptomatic case (blue) at the ends of major outbreaks (outbreaks in which more than 20 individuals are ever infected). The current WHO guideline period of 42 days leads to a confidence of 0.84 that the outbreak is over (red), whereas periods of 62 or 88 days correspond to confidences of 0.95 (light green) and 0.99 (dark green), respectively. E) The confidence in end-of-outbreak declarations made 42 days after removal of the “final” symptomatic case, for different values of the surveillance sensitivity (blue). For an end-of-outbreak confidence of 0.95 after 42 days without symptomatic cases, a surveillance sensitivity of 79% is required (light green). The results in panels C and D were obtained using 10,000 simulations of the model, and the results in panel E were obtained using 10,000 simulations of the model for each possible value of the surveillance sensitivity.

The proportion of cases identified by public health authorities through passive or active case finding, otherwise called the sensitivity of a surveillance system (*6,7*), is a critical parameter that underlies how confident a decision-maker can be when declaring the end to an outbreak (*8*). The surveillance sensitivity is the ratio of the number of infectious cases detected to the total number of cases (including both cases that are detected and those that go unnoticed), and should not be confused with the sensitivity of a diagnostic test (i.e. the probability that the diagnostic test correctly identifies an infected host). Intuitively, there will be a lower confidence that an outbreak is over if the surveillance system has low sensitivity. However, decision-makers do not typically make quantitative assessments about the confidence in their end-of-outbreak decisions. A retrospective modelling study of the MERS-CoV outbreak in South Korea in 2015 concluded that, with no quantitative end-of-outbreak assessment, decision-makers took longer than epidemiologically necessary to declare the outbreak over (*9*), although we note that the sensitivity of the surveillance system was assumed to be 100% in that study.

To illustrate how the confidence that an outbreak is over can be estimated, and that the confidence level can be increased by improving surveillance, we consider the situation of declaring the end of an Ebola outbreak. The WHO considers an Ebola outbreak to be over once 42 days have passed since the last possible exposure to a confirmed case without any new cases being detected (*10*), with this rule most often deployed at the scale of a single country. The incubation period (the time between an individual becoming infected and displaying recognisable symptoms) for Ebola has been estimated to be in the range of 2-21 days (*11*), and so the period of 42 days is based on two maximal incubation periods. For a disease that is transmitted directly from person-to-person, the passing of two incubation periods is epidemiologically relevant because additional between-person transmission is then unlikely. We use mathematical modelling to show that, if the surveillance sensitivity is 100%, then it is likely that an Ebola outbreak is over after 42 days without symptomatic cases. However, we also demonstrate that, if the surveillance sensitivity is lower, there is an increased chance of undetected infected cases remaining after 42 days. This leads to a lower confidence that the outbreak is over after this time period.

Although we focus on Ebola virus disease here, we note that the question of whether or not an infectious disease outbreak is over is not only important for diseases of humans, but also those of animals and plants. Declaring an outbreak over allows disease management interventions to be lifted, including restrictions on travel (*12*) and plant trade quarantine (*13*). The idea that improved surveillance may lead to increased confidence in an end-of-outbreak declaration is related to well-established theory regarding conducting surveys to ascertain the absence of a pathogen (see e.g. (*14-19*)). In that context, the more hosts are tested and found to be disease-free, the higher the confidence that the entire population is disease-free. This can in turn be used to generate sample-size requirements to establish freedom from disease to pre-specified confidence levels. While initial studies in this area – motivated by the desire to limit pathogen transmission via the animal trade – assumed that the level of disease in the host population was static, more recent elaborations have included incorporation of dynamic models describing parasite/pathogen transmission in the host population (see e.g. (*20,21*)). Statistical disease freedom studies have not only been applied to animal disease epidemics, but the theory has also been used in the context of epidemics in populations of plants (*22, 23*) and humans (*24*).

In this paper, rather than considering surveys of the host population at the apparent end of an outbreak, we show how the confidence in end-of-outbreak assessments can be estimated using epidemiological models once the surveillance system sensitivity has been approximated. Our approach, which can be used when the outbreak in question is still ongoing, provides decision-makers with a practical way to gauge whether current surveillance efforts will be sufficient to declare the outbreak over with conviction, or whether an intensification of surveillance is necessary instead.

## Methods

We extended an epidemiological model commonly used for Ebola (the SEIR model, see e.g. (*25,26*)) to include imperfect surveillance (Fig 1B). In the resulting model (the SEICR model) individuals were classified according to whether they were (*S*)usceptible, (*E*)xposed, (*I*)nfectious and reporting disease, (*C*)ryptically infectious (i.e. infectious but not reporting disease), or (*R*)emoved. The parameters of the model and the baseline values used in our analyses to illustrate the model behaviour are given in Table 1, although we also tested the robustness of our results to these values (Fig S1). We ran stochastic simulations of the model, thereby including randomness in whether or not each outbreak was over when the number of symptomatic individuals (*I*) reached zero (for additional details, see the Supplementary Material).

**Table 1.** Epidemiological parameters of the SEICR model, along with the default values used in our analyses (except where stated in the relevant figure captions) and references supporting the values used. We also tested the robustness of our results to the values of *N, d* and *β* (Fig S1). For more details about the model and its parameterisation, see Supplementary Material.

| <u>Epidemiological parameter</u> | <u>Meaning</u> | <u>Baseline value (used except where stated)</u> | <u>Justification</u> |
| --- | --- | --- | --- |
| $\beta$ | Infection rate | $2.7 \times 10^{-6} \text{ day}^{-1}$ | Chosen such that $R_0 = 2$ (see e.g. (46,47)) |
| $1/\gamma$ | Incubation period | 12.27 days | (48) |
| $1/\mu$ | Infectious period | 7.37 days | (48) |
| $N$ | Effective population size | 100,000 | For consistency of mean final size of simulated epidemics with 2013-16 Ebola epidemic |
| $d$ | Proportion of infectious hosts reporting disease | 0.4 | (28) |

The surveillance sensitivity was implemented in the model via the proportion, *d*, of infectious individuals that reported disease (*I*) as opposed to remained cryptically infectious (*C*). When an individual left the exposed class, they either transitioned into the *I* class (with probability *d*) or into the *C* class (with probability 1 – *d*). The parameter *d* represents a proportion/probability and therefore lies between zero and one, whereas the surveillance sensitivity is reported as a percentage. As an example, the value *d* = 0.1 corresponds to a surveillance sensitivity of 10%. For simplicity, we assumed that whether or not an infectious individual reported disease did not alter their infectiousness or duration of infection, although this simplification could be relaxed straightforwardly. As described above, the cryptically infectious class represents individuals that are infectious but do not report disease – this could include asymptomatic carriers that are infectious (*27*) or symptomatic individuals not reporting for reasons including a lack of access to healthcare (*8*).

By continuing to run simulations after the number of symptomatic infectious individuals (*I*) reached zero, the confidence that an outbreak will be over, defined as the probability that no undetected infected hosts (*E* or *C*) remained in the population, was estimated at different time periods beyond the removal of the last detected case.

## Results

We inferred the expected number of undetected infected cases once the number of symptomatic cases reached zero (Fig 1C), considering only outbreaks that successfully invaded the host population (outbreaks in which more than 20 individuals were ever infected). We estimated the confidence that the outbreak is over for different time periods beyond the removal of the last detected case (Fig 1D). For additional results with different model parameters, see the Supplementary Material.

Since a period of 42 days has been estimated as twice the maximal incubation period for Ebola, it is unsurprising that, when the sensitivity of the surveillance system was perfect so that 100% of infectious cases were detected accurately, the model suggested a high confidence (more than 97%) that an Ebola outbreak is over after 42 days without symptomatic cases. Additional new cases could only occur if existing infected individuals remained pre-symptomatic, and this was very unlikely after this time period. However, when we assumed that the surveillance sensitivity was only 40%, an estimate made for Ebola surveillance in Liberia (*28*), the probability that Ebola cases remained in the population was 16%, leading to only an 84% confidence that the outbreak was finished (red line in Fig 1D). With such a low surveillance sensitivity, a period of 62 days with no cases would need to elapse to be 95% confident that an outbreak is over (light green line in Fig 1D), or 88 days to be 99% confident (dark green line in Fig 1D).

Most Ebola cases in outbreak areas are reported via infected individuals presenting to a health facility or Ebola Treatment Unit. Close contacts of confirmed cases are also identified and usually followed for 21 days, to permit rapid identification if symptoms develop (*29*). Other case finding strategies may take place, for example visitations to identify and test suspect cases in disease hotspot regions or in rural areas where access to healthcare systems might be limited (*30*). Since surveillance can potentially be improved, for example by intensifying active case finding, the quantity of most practical value is the sensitivity of surveillance required for decision-makers to be confident that Ebola outbreaks are over after 42 days. We therefore considered the end-of-outbreak confidence for varying levels of the surveillance sensitivity (Fig 1E). To be at least 95% confident that an Ebola outbreak is over after 42 days, surveillance needed to be at least 79% sensitive (light green line in Fig 1E). For lower surveillance levels, there is a significant chance (> 5%) of residual infectious cases remaining in the population, and these might generate outbreak flare-ups.

## Discussion

We have proposed an approach for decision-makers to estimate their confidence that an Ebola outbreak is over after 42 days (two maximal incubation periods) have passed with no new cases. In scenarios with a low surveillance sensitivity, decision-makers may either choose to wait longer than two incubation periods before declaring the end of an outbreak, take measures to increase the surveillance sensitivity, or adopt both of these approaches. Communicating that an outbreak is over following two incubation periods is epidemiologically coherent when the surveillance level is high, and so decision-makers may prefer to focus efforts on achieving a high surveillance sensitivity rather than adjusting the guideline period before declaring the end of an outbreak. However, in contexts that prevent strengthening of disease surveillance, for example if there is poor security due to armed conflict or other factors, extending the period with no cases before declaring an outbreak over may be the more pragmatic option.

Sensitivity measurements are sometimes carried out for evaluation of surveillance systems (*31*). Analysis of the percentage of cases being recorded can be conducted using serological surveys (*32*) or by comparing multiple data sources (*33*). When an outbreak is ongoing, however, measuring the surveillance sensitivity might not be the first priority. For assessing the confidence in a potential end-of-outbreak declaration, it is most important to measure the surveillance sensitivity towards the apparent end of the outbreak, and so resources can be directed to this task after the acute outbreak period has passed. In scenarios in which the surveillance sensitivity is insufficient for declaring an outbreak over with confidence, remedial actions can be taken such as strengthening case finding for example via contact tracing (*34*), closer working with community leaderships to establish a case finding and reporting network (*35*), and/or providing incentives for successful case reporting (*36*), among other approaches.

In this paper, we sought to use a simple approach to demonstrate how the confidence in end-of-outbreak declarations could be assessed, and to show that rigorous surveillance is extremely important. While accurate case reporting will minimise the chance of incorrect declarations that Ebola outbreaks are over in future, we note that surveillance during the outbreak alone is not always sufficient. In the 2013-16 Ebola epidemic in West Africa, additional cases occurred after regions were declared disease-free due to factors including persistently infected sources (*37*) and importation of the virus from other geographical regions (*38*). There were suspicions of a flare-up arising from a female survivor, who became infectious after her immune system was weakened due to pregnancy (*39*), although this remains unproven (*40*). There is also evidence that Ebola survivors might have the potential to drive new cases after long periods following apparent recovery (*41*), for example reports of the virus being detected in semen up to 18 months after symptom onset or isolated in cell culture up to 82 days after symptom onset (*42,43*). Here, we only considered potential flare-ups due to unreported cases, and did not explicitly model the possibility that Ebola survivors, who were assumed to have fully recovered, might drive additional cases. Recrudescence from survivors could, in theory, be included in assessments of the risk of outbreak flare-ups after outbreaks are declared over, however this would require sufficient understanding of the epidemiology of these rare events. While this understanding is developed, targeted monitoring of survivors beyond the WHO guideline period of 42 days that we consider here is also important. This should be supplemented with advice for survivors on safe practices that will help to avoid additional flare-ups (*44*).

Extending our approach to other disease outbreaks might require elaborations to the underlying model. To illustrate the principle that the surveillance sensitivity affects the confidence in end-of-outbreak declarations, we modelled surveillance as simply as possible – by assuming that a proportion of infectious hosts report disease, but that reporting did not impact on the underlying transmission process. In practice, individuals that report disease are more likely to be subject to interventions that reduce infectiousness or shorten their infectious period, such as isolation or treatment. This could straightforwardly be built into the framework that we have presented. We also use a single parameter to denote the surveillance sensitivity, whereas in practice a surveillance program is likely to encompass many aspects, including both passive and active case finding strategies, that could be built explicitly into an epidemiological model. One of the benefits of our approach is that, in contrast to methods relying on surveys to prove disease absence, our analysis can be conducted in advance of the apparent end of the outbreak to see whether or not surveillance needs to be intensified. However, it might be possible to combine our approach with surveys to establish the end of an outbreak, and to make use of statistical methods for estimating the number of hosts to survey so that the probability of the population being disease-free exceeds a pre-specified threshold (*14-24*).

Other extensions could include modelling the risk of importation of disease from other geographical locations (*13*), accounting for temporal or spatial variation in the surveillance sensitivity (*8*), or allowing for the possibility of introductions of immunologically naïve hosts resulting from population displacement (*45*). Additional refinement would be needed to estimate the confidence in end-of-outbreak declarations when diseases that persist at low endemic levels in a population have returned below an outbreak threshold prevalence, an important consideration for outbreaks of diseases such as cholera and yellow fever. Nonetheless, we have demonstrated that measurement of the sensitivity of surveillance can help decision-makers estimate their confidence that an outbreak has ended. Moreover, after measuring the performance of disease surveillance systems, the confidence that an outbreak has ended can be increased by optimising the surveillance system.

Communicating the confidence in end-of-outbreak decisions, as our framework permits, is helpful for decision-makers and the public, and it will increase trust in public health organisations such as WHO. Measurements of the surveillance sensitivity during outbreaks are not always routinely taken, but we think that by providing a useful way to use measures of surveillance system performance, decision-makers will be motivated to implement what should be considered as good practice. We encourage the use of quantitative approaches, such as the one we describe here, to inform decisions regarding the continuation of disease control measures, appropriate use of resources, and communication of public health messages to the public towards the end of an outbreak.

## Supporting information

## Acknowledgments

Thanks to the organisers of the 2018 Hackout meeting, particularly Thibaut Jombart, at which discussions about this work took place between RNT, OWM and KJ. Thanks to Amy Dighe and Finlay Campbell for helpful comments.

## Authors’ contributions

RNT conceived the research; All authors designed the study; RNT carried out the research; RNT drafted the manuscript; All authors revised the manuscript and gave final approval for publication.

