## Supplementary material for "Rigorous surveillance is necessary for high confidence in end-of-outbreak declarations for Ebola and other infectious diseases"

**Epidemiological model**

We performed our analyses using a stochastic compartmental model characterising an Ebola virus disease outbreak. The classic deterministic SEIR model has the following form:

$$\frac{dS}{dt}=-\beta SI,$$

$$\frac{dE}{dt}=\beta SI-\gamma E,$$

$$\frac{dI}{dt}=\gamma E-\mu I,$$

$$\frac{dR}{dt}=\mu I.$$

We extended this model to include a new class (*C*), consisting of infected individuals who do not report their disease, leading to the SEICR model:

$$\frac{dS}{dt}=-\beta S\left( I+C \right),$$

$$\frac{dE}{dt}=\beta S(I+C)-\gamma E,$$

$$\frac{dI}{dt}=\gamma dE-\mu I,$$

$$\frac{dC}{dt}=\gamma\left( 1-d \right)E-\mu C,$$

$$\frac{dR}{dt}=\mu(I+C).$$

The key difference between the SEICR model and the SEIR model is that, in the SEICR model, only some infectious individuals – a proportion *d* – are assumed to report disease (see Model parameters, below). As a result, the outbreak is not necessarily over when the final symptomatic infectious host is removed from the *I* class. We performed simulations of the analogous stochastic SEICR model, using the Gillespie direct method (Gillespie *et al*., 1977) to determine precise transition times of individuals between compartments. Most of our model simulations were run in an effective host population size of *N* = 100,000, starting from a single symptomatic infected individual (*I* = 1) with the remainder of the population susceptible (*S* = *N* - 1), although the choice of effective population size was not central to our results (Fig S1A).

**Model parameters**

The following default parameter values were used, except where stated. The mean incubation period for symptomatic infected individuals was $1/\gamma$ = 12.27 days (Velásquez *et al*., 2015), and for simplicity this quantity was assumed to be identical to the latent period, although we note that there has been some debate as to whether or not infectiousness and symptoms always coincide (Velásquez *et al*., 2015) and the extent to which modellers should account for any differences (Thompson and Hart, 2018). The infectious period was $1/\mu$ = 7.37 days. The default proportion of infectious individuals assumed to report disease was $d$ = 0.4 (i.e. a surveillance sensitivity of 40%), in line with estimates from the World Health Organization (Meltzer *et al*., 2014), although we varied this parameter in different analyses (see Fig 1E of the main text and Fig S1B). An infection rate parameter of $\beta=2.7\times{10}^{-6}$ per day was used, yielding a basic reproduction number of 2, consistent with published estimates for Ebola (see e.g. Van Kerkhove *et al*., 2015; Krauer *et al.*, 2016). These parameters are also summarised in Table 1 of the main text.

**Robustness of results**

By using a standard compartmental modelling approach in our analyses, we assumed a well-mixed population, with any individual having an equal chance of contact with any other (Keeling and Rohani, 2007). As a result, we assumed a smaller effective host population size of *N* = 100,000 than that of an entire city or country, in which the population is not well-mixed. The mean final size of major outbreaks (outbreaks for which the final value of *R* was greater than 20) simulated with our default parameters was 79,505, consistent with the order of magnitude of the total number of cases in the 2013-16 Ebola epidemic in West Africa adjusted for under-reporting. However, we also tested the robustness of our results to the value of the effective population size *N* (Fig S1A), as well as to the surveillance sensitivity (Fig S1B) and the basic reproduction number (Fig S1C).

In each of the different scenarios shown in Fig S1 for which the surveillance sensitivity was less than 79%, there was a significant chance (> 5%) that undetected, infected cases remained in the population 42 days after the “final” symptomatic individual was removed. By increasing the sensitivity of surveillance, governed by the case reporting parameter *d*, the confidence in end-of-outbreak declarations will be increased (Fig S1B and Fig 1E in main text). This motivates our recommendation that, for a confidence level of 95% that outbreaks are really finished when declared over, the surveillance sensitivity should be increased to above 79% via active case finding at the tail-ends of Ebola outbreaks.

By applying our model to data collected during an ongoing outbreak, it would also be possible to condition the estimate of the confidence in an end-of-outbreak declaration to data from that specific outbreak. Developing methods for this purpose would make an interesting target for further study, although in practice general recommendations for all outbreaks of a particular disease like the one proposed here – to aim to increase the surveillance sensitivity to above 79% during Ebola outbreaks by active case finding – might be more relevant for practical application.

**Supplementary figure**

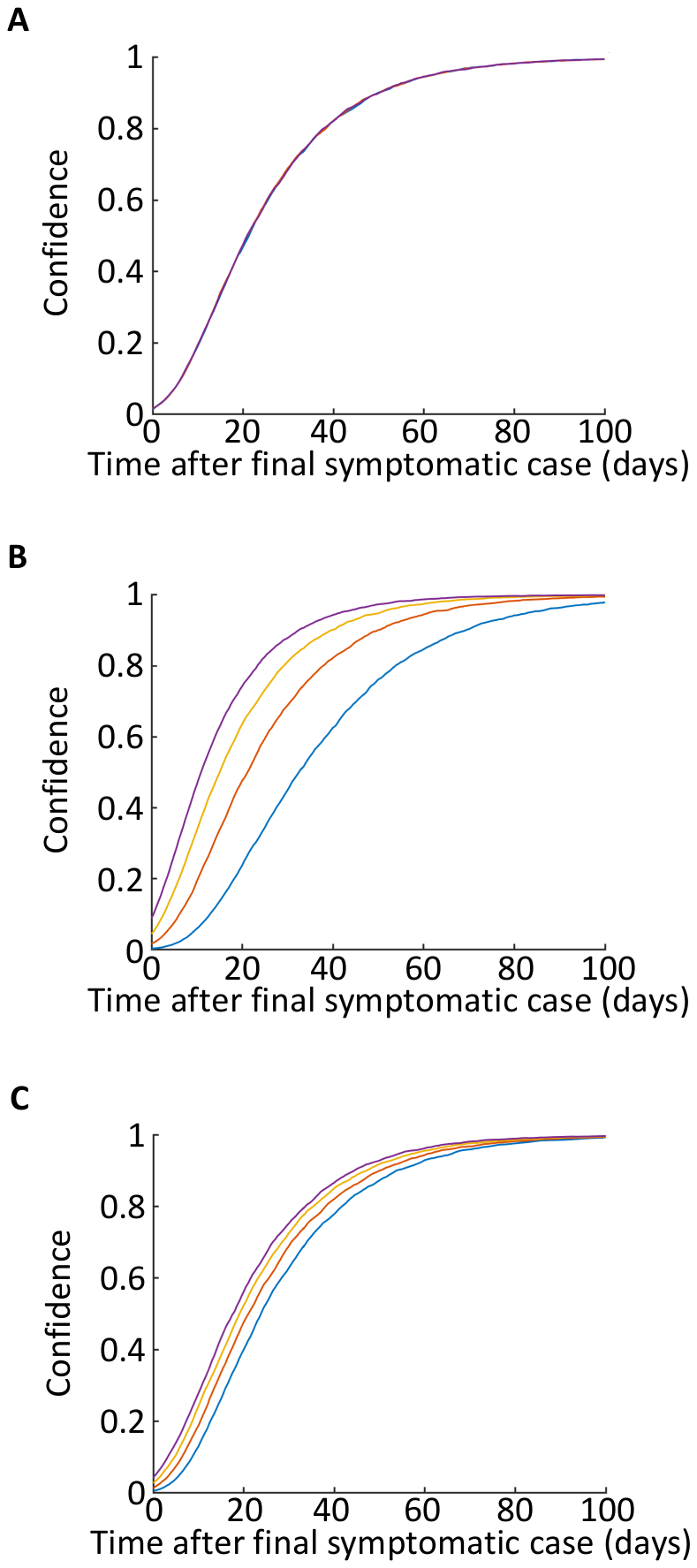

Figure S1. The confidence in end-of-outbreak declarations at different times following the apparent end of the outbreak, for a range of model parameters. A) Different values of the effective population size, *N* (blue – *N* = 1,000; red – *N* = 10,000; yellow – *N* = 100,000; purple – *N* = 1,000,000). B) Different values of the reporting fraction, *d* (blue – *d* = 0.2 (i.e. surveillance sensitivity of 20%); red – *d* = 0.4 (surveillance sensitivity of 40%); yellow – *d* = 0.6 (surveillance sensitivity of 60%); purple – *d* = 0.8 (surveillance sensitivity of 80%)). C) Different values of the basic reproduction number, *R*_0_, varied by changing $\beta$ (blue – *R*_0_ = 1.2; red – *R*_0_ = 1.6; yellow – *R*_0_ = 2; purple – *R*_0_ = 2.4). Default parameter values for *N*, *d* and *R*_0_ were used except where stated, so that e.g. in panel A, *N* is varied while holding *d* and *R*_0_ constant. In particular, in panel A, when *N* was varied then $\beta$ was also varied to maintain a constant value of *R*_0_ = 2. These results were obtained using 10,000 simulations of the model per line in each panel.
